# A comparative *in vitro* and *in vivo* study of osteogenicity by using two biomaterials and two human mesenchymal stem cell subtypes

**DOI:** 10.1101/2021.07.17.452782

**Authors:** Lucile Fievet, Nicolas Serratrice, Bénédicte Brulin, Laurent Giraudo, Julie Véran, Nathalie Degardin, Florence Sabatier, François Féron, Pierre Layrolle

## Abstract

Bone repair induced by stem cells and biomaterials may represent an alternative to autologous bone grafting. Here, we compared the efficiency of two biomaterials - biphasic calcium phosphate (BCP) and bioactive glass (BG) - when loaded with either adult bone marrow mesenchymal stem cells (BM-MSCs) or newborn nasal ecto-mesenchymal stem cells (NE-MSCs), the latter being collected for further repair of lip cleft-associated bone loss. Both cell types display the typical stem cell surface markers CD73+/CD90+/CD105+/nestin, and exhibit the MSC-associated osteogenic, chondrogenic and adipogenic multipotency. NE-MSCs produce less collagen and alkaline phosphatase than BM-MSCs. At the transcript level, NE-MSCs express more abundantly three genes coding for bone sialoprotein, osteocalcin and osteopontin, while BM-MSCs produce extra copies of *RUNX2*. BM-MSCs and NE-MSCs adhere and survive on BCP and BG. *In vivo* experiments reveal that bone formation is only observed with BM-MSCs transplanted on BCP biomaterial.

## Introduction

Biomaterial- and stem cell-associated bone repair represents a very promising alternative to autologous bone grafting. The most commonly investigated cells are mesenchymal stem cells (MSCs)^**1–4**^, among them, bone marrow derived MSCs (BM-MSCs) tend to be considered as the gold standard due to their capacity in combination with biomaterial scaffolds to repair bones *in vivo*^5–7^. However, other sources of MSCs have been poorly investigated^**8**^. Our laboratory identified and purified from the *lamina propria* of the human olfactory mucosa, the ecto-mesenchymal stem cells, originate from the neural crest, that reside in the craniofacial area during adulthood^**9–11**^. When compared to BM-MSCs, nasal ecto-mesenchymal stem cells (NE-MSCs) display an inclination to differentiate into osteocytes, rather than chondrocytes or adipocytes, and produce mineralized bone tissue *in vivo* after being filled on calcium phosphate ceramic discs when implanted subcutaneously in nude mice ^**12**^.

Many biomaterials have been developed to offer surgeons and patients safe alternatives to autologous bone graft, that is limited in quantity and adds morbidity at the harvesting site. The most commonly used are composed of biphasic calcium phosphate (BCP), a mixture of hydroxyapatite (HA)/beta-tricalcium phosphate (βTCP), similar to the mineral bone composition, but they display an insufficient osteoinductive capacity to regenerate large bone defects. BM-MSCs added to porous calcium phosphate ceramics induced bone formation when implanted under the skin of nude mice^**6,13–15**^, in rat and pig femurs^**16,17**^. This BCP/BM-MSCs combination has even demonstrated safe bone regeneration in clinical trials^**18–20**^.

Most recently, a synthetic bioactive glass (BG) or bioglass (bioglass 45S5 or GlassBONE^TM^) has also been used for bone regeneration in orthopedic, traumatic, spinal and craniomaxillo-facial surgery. Based on borate and borosilicate, BG enhances bone and underlying neocartilage formation when compared to silicate bioactive glass^**21,22**^. Moreover, borate-based BG displays controllable degradation rates, and can be doped with trace quantities of elements such as Cu, Zn and Sr, which are known to be beneficial for healthy bone growth. In addition, it has been observed that BG promotes angiogenesis, which is critical in bone tissue regeneration^**23**^. In order to go further, we design a new study comparing the osteogenic capacity of human mesenchymal stem cells (hMSCs), derived from either adult bone marrow or newborn nasal cavity (having in the mind the repair of cleft lips), associated with BCP or BG biomaterials, *in vitro* and after *subcutis* implantation in nude mice.

## Results

### Newborn nasal stem cells display an NE-MSC phenotype

Human newborn nasal cells being cultivated for the first time in the team, we first assessed their stemness. They express nestin (**Figure 1A**), a widely recognized marker for stem cells. In addition, they produce CD73, CD90 and CD105 (**Figures 1B-D**), as documented for adult NE-MSCs, and not CD45, a specific hematopoietic stem cell marker (**Figure 1E**).

**Figure 1:**
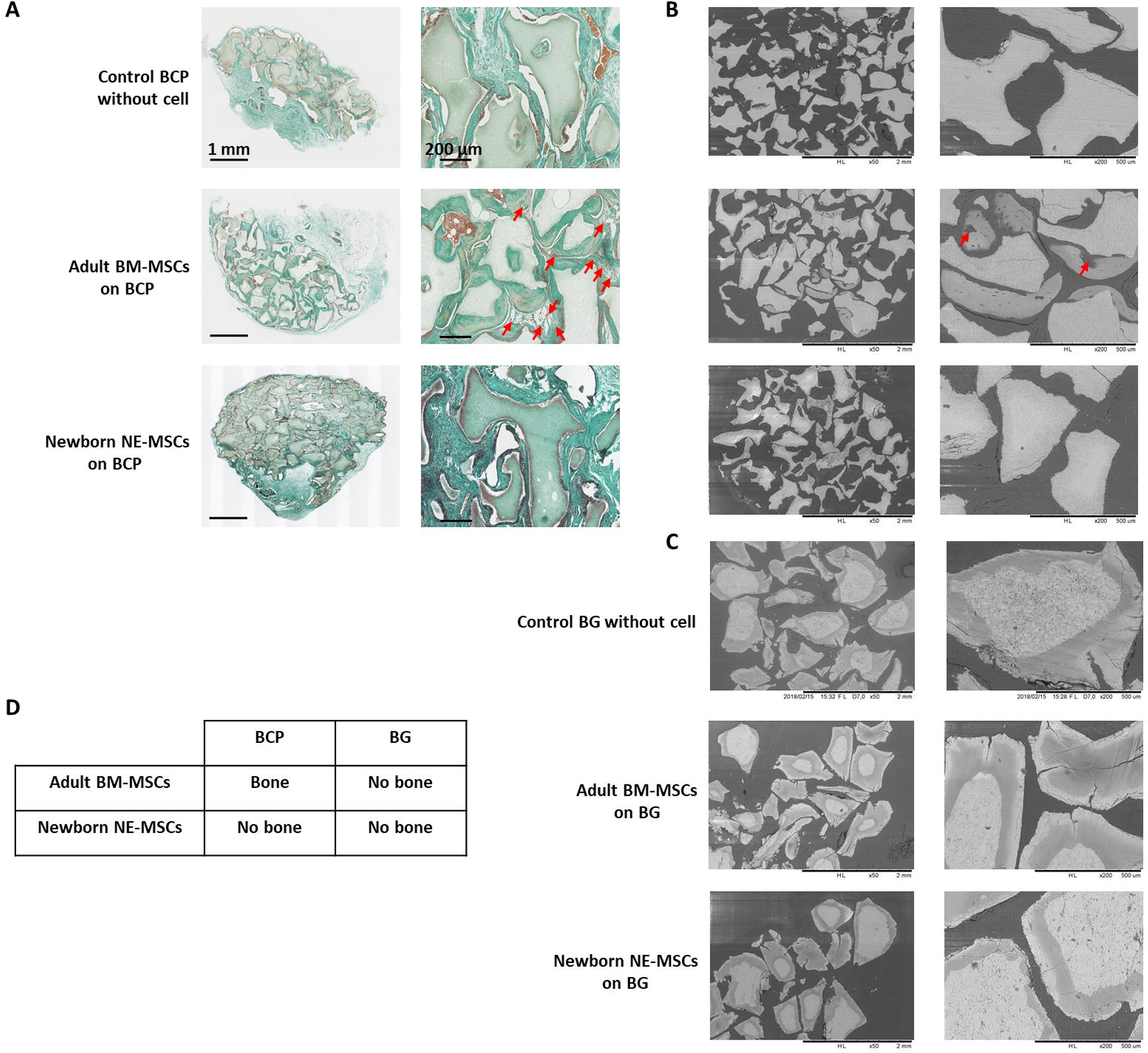
Characterization of newborn nasal stem cells. Phenotyping of cultivated cells was performed using flow cytometry and stem cell-specific markers. Purified nasal cells were positive for nestin (**A**), CD73 (**B**), CD90 (**C**) and CD105 (**D**), recognized markers of NE-MSCs, and negative for CD45 (**E**), known marker for hematopoietic stem cells. 10,000 events counted.

### In vitro assessment of stem cell multipotency

We compared the capacity of both MSC types to differentiate *in vitro* into osteocytes, chondrocytes, and adipocytes, using the usual stains: alizarin red, alcian blue, and red oil (**Figure 2A-C**). BM-MSCs and NE-MSCs display an intense mineralization at D14, and exhibit a similar pattern of differentiation for chondrocytes and adipocytes.

**Figure 2:**
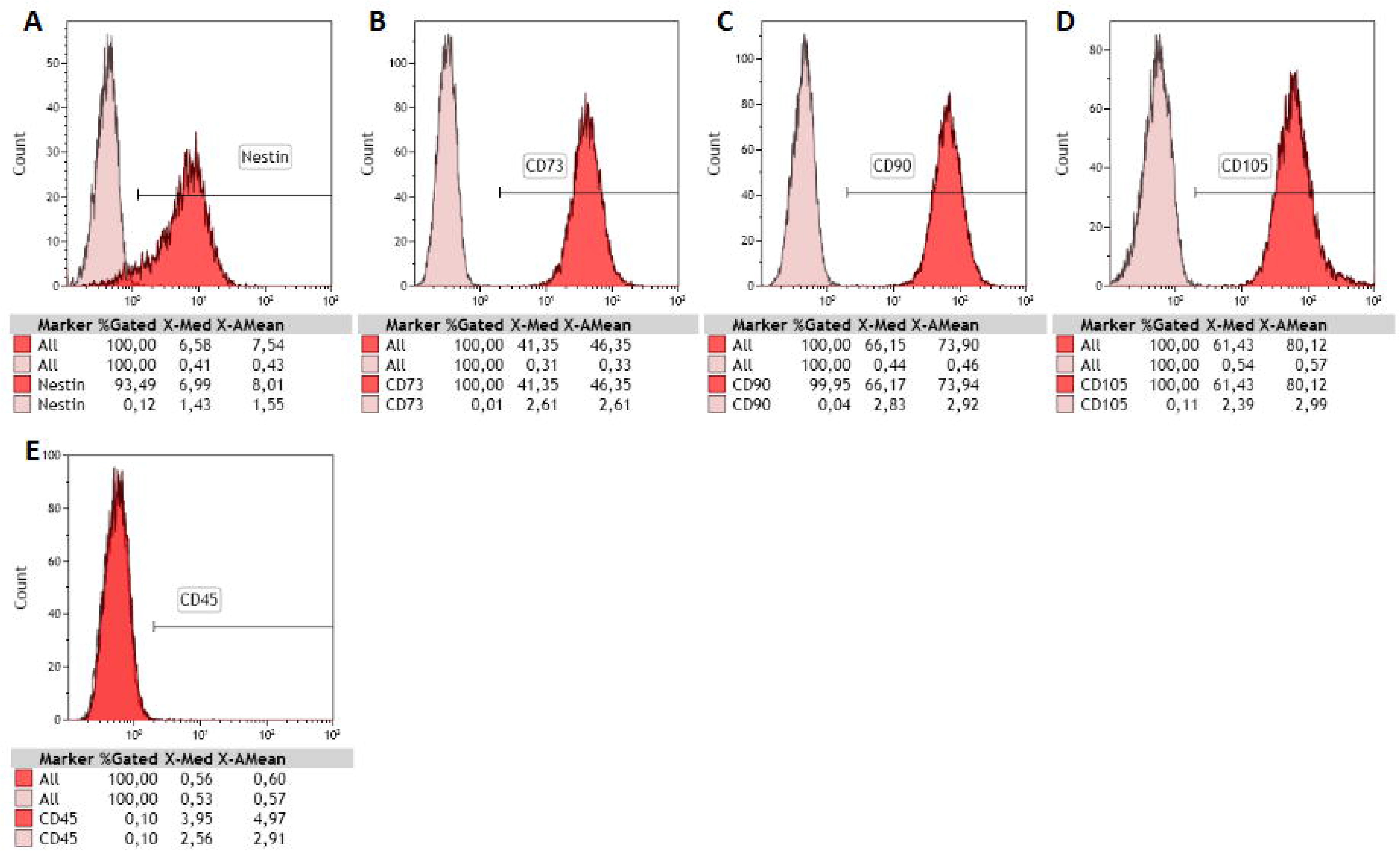
*In vitro* assessment of multipotency. BM-MSCs and NE-MSCs were induced to differentiate into osteocytes (alizarin red, **A**), chondrocytes (alcian blue, **B**) and adipocytes (red oil, **C**). Both BM-MSCs and NE-MSCs showed mineralization production at D14. NE-MSCs differentiated in chondrocyte at D14, while BM-MSCs at D21. BM-MSCs and NE-MSCs also differentiated in adipocytes at D14.

### BM-MSCs and NE-MSCs adhere and survive on biomaterials

Both BM-MSCs and NE-MSCs adhere to BCP and BG (**Figure 3A**). BG, and not BCP, induces cell death (**Figure 3B**), as early as D2 for BM-MSCs, and only at D21 for NE-MSCs, a possible indication that stem cells from human newborns are more resistant to a relatively adverse material.

**Figure 3:**
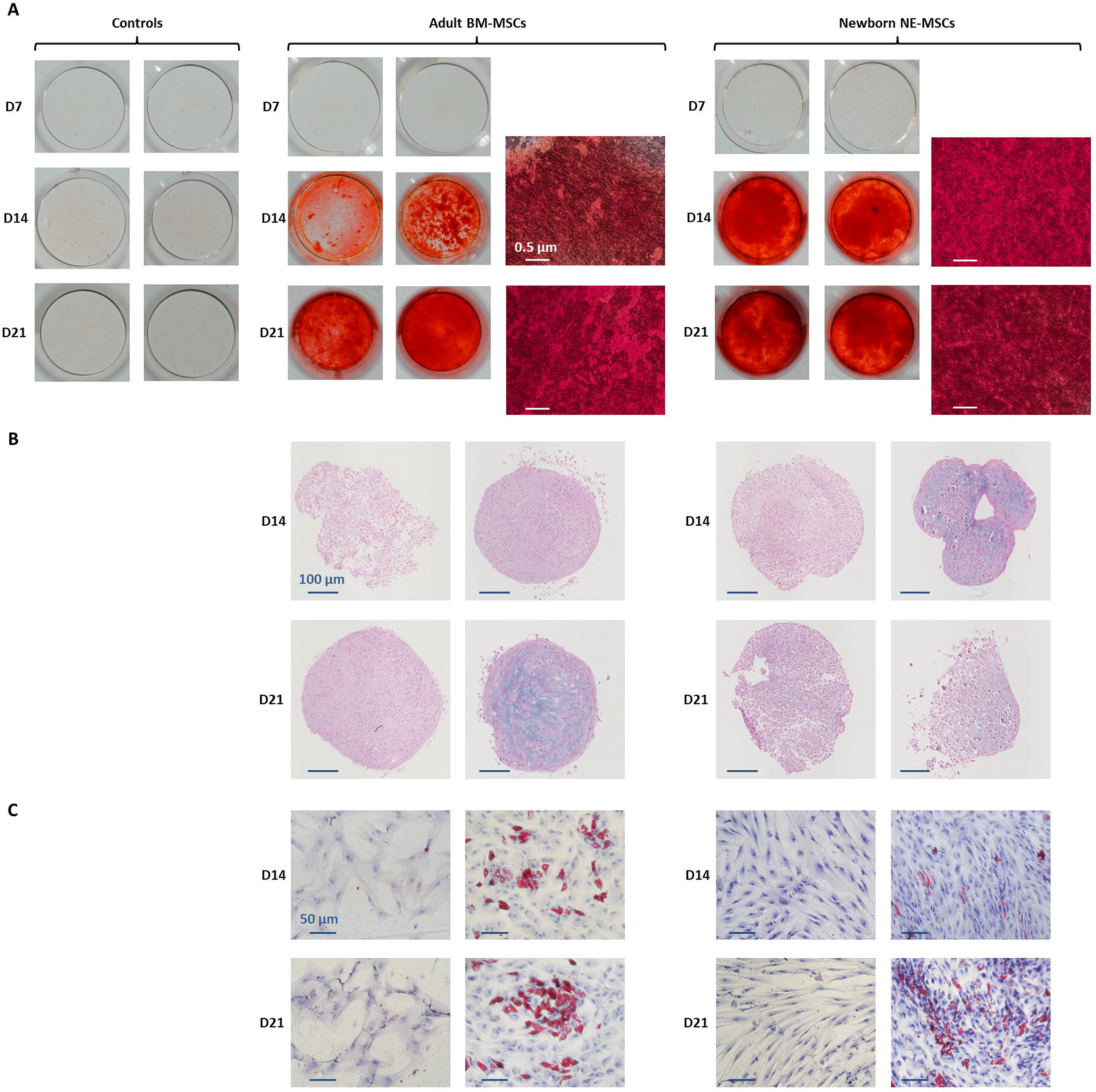
(**A**) BCP and BG 3D constructs visualization by SEM at D2, D7, D14, and D21. BCP and BG had similar shape and size but different surface microstructures, BCP particles showed microporosity, whereas BG exhibited a smooth surface. Both types of MSCs adhered more rapidly to BG than to BCP. BM-MSCs synthesized an extracellular matrix after D14, which seem to be more abundant on BG than on BCP, but mineralization was only observed on BCP. NE-MSCs produced also an abundant, but fragile, mineralized extracellular matrix on both biomaterials (less on BG than on BCP). (**B**) Comparative adhesion and viability of the two MSC types on biomaterials. Using fluorescent stains for living (green) and dead (red) cells, BM-MSCs and NE-MSCs viability was kinematically assessed. Dead cells (white arrows) are observed only on BCP biomaterial. Death of newborn stem cells was only observed at D21 while apoptosis/necrosis of adult stem cells starts as soon as D2.

### BM-MSCs display a higher proliferation on BG biomaterial

On plastic, BM-MSCs and NE-MSCs display a high proliferation rate, although the latter overcomes the former^12^. It was then questionable to confirm this finding when using biomaterials. **Figure 4** indicates that, on both biomaterials, NE-MSCs proliferate more rapidly than BM-MSCs at the start of the experiment. However, at D14, the numbers are even and, subsequently, due to cell death occurring at this stage (see above), the density of both MSCs declines (**Figures 4A-B**). **Figures 4C-E** indicates that, when compared to BCP, BG favors BM-MSC proliferation. No difference between the two materials observed for NE-MSCs.

**Figure 4:**
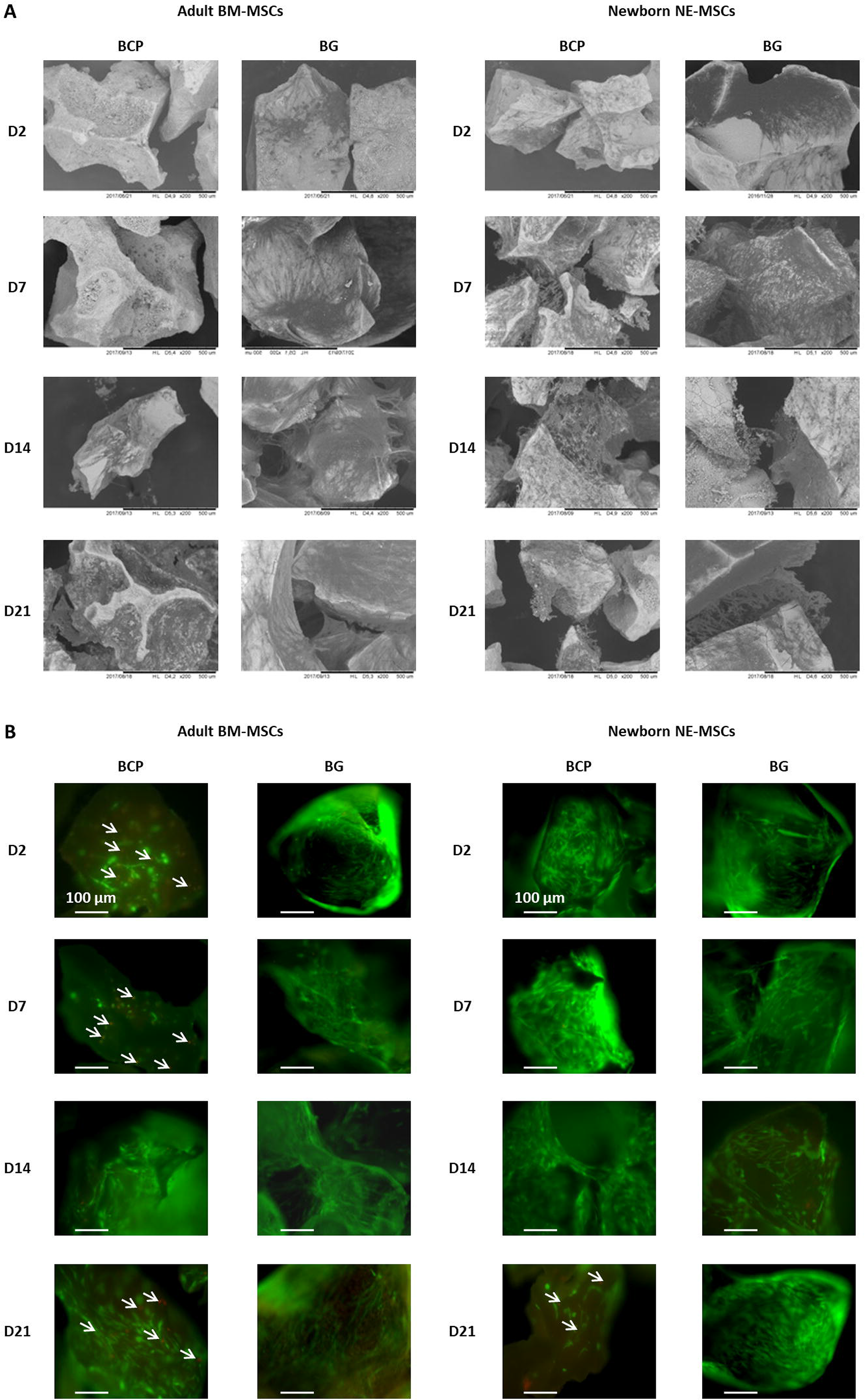
Influence of the biomaterial on cell proliferation. BM-MSCs and NE-MSCs were seeded on 2D plastic at two different densities: 500 (**A**) and 5,000 cell/cm^2^ (**B**). (**C-E**) Both cell types were seeded on BCP and BP and cell metabolic activity monitored over a period of three weeks using alamarBlue^®^ assay. On plastic, NE-MSC density increases more rapidly during the first week but an equalization occurs at the end of the second week. During the third week, cell death induces a reduced confluence. **C-E graphs** indicate that, when compared to BCP, BG biomaterial favors BM-MSC proliferation. Controls corresponds to cell-free biomaterials (background noise of alamar blue).

### Collagen production increases in BM-MSCs and decreases in NE-MSCs

BM-MSCs produce more collagen, being a chondrocytic and osteogenic specific marker (Sirius red staining), than NE-MSCs. Noticeably, collagen expression timely increases in BM-MSCs, and declines in NE-MSCs (**Figures 5A**).

**Figure 5:**
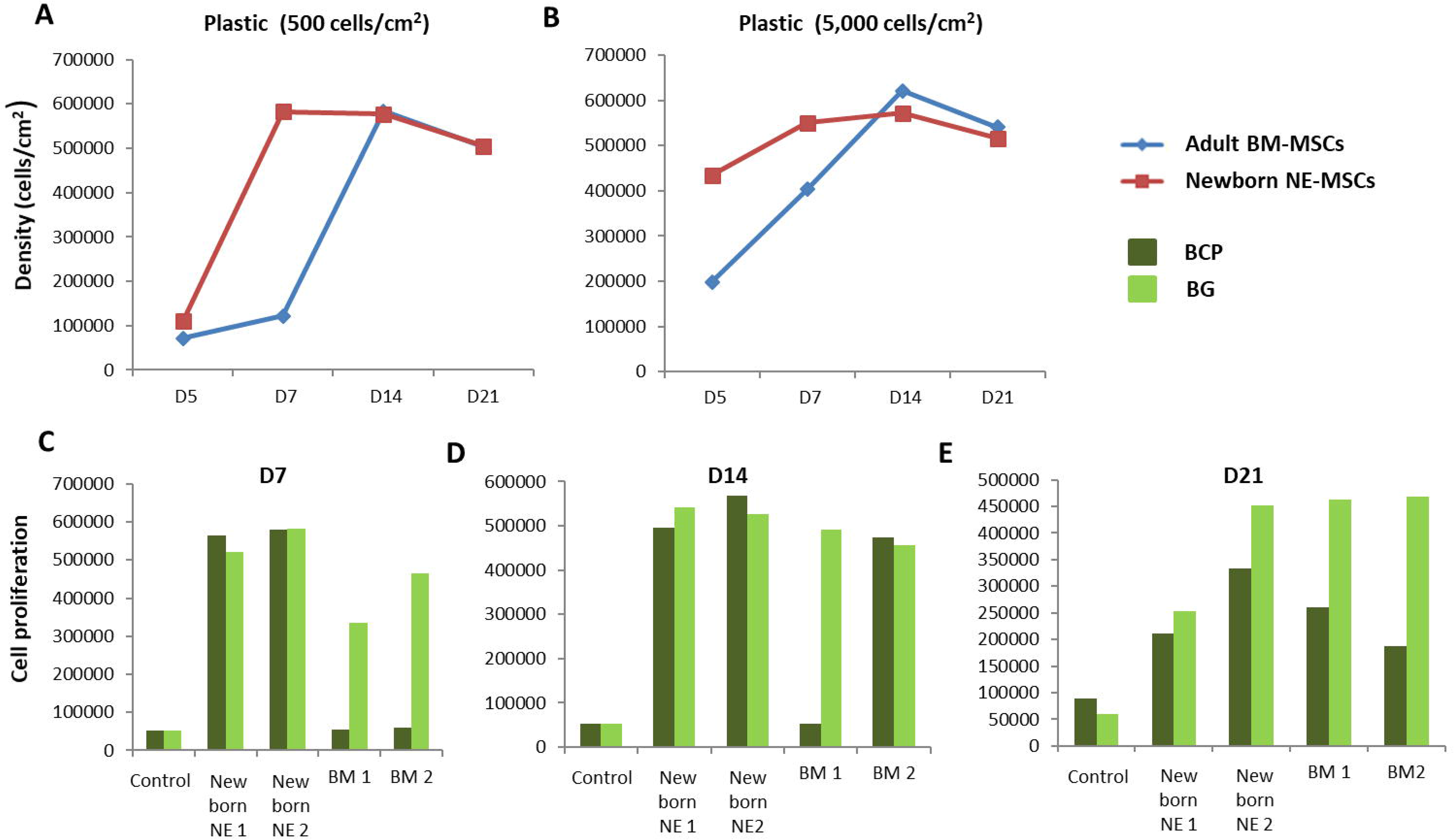
(**A**) Comparative production of collagen. In osteogenic conditions, BM-MSCs increasingly produce collagen while the opposite is observed for NE-MSCs. Comparative production of ALP by both cell types, at D7. BM-MSCs and NE-MSCs were cultivated either on plastic (**B**) or biomaterials (**C**). BM-MSCs express ALP only when cultivated on BG biomaterial while NE-MSCs produce ALP on both biomaterials. For NE-MSCs, ALP production is higher in proliferative medium.

### BM-MSCs produce ALP only on BG biomaterial

BM-MSCs and NE-MSCs produced ALP on plastic (**Figure 5B**). At D7, BM-MSCs express ALP only when cultivated on BG, while NE-MSCs produce ALP on both biomaterials with a higher production in proliferative medium (**Figure 5C**). To move further and quantify ALP production in various conditions, we assessed its expression in accordance with cell abundance (**Figure 6A-B**). NE-MSCs express an important amount of *ALP* when grown on BG in osteogenic conditions, and conversely is negligible on BCP, except at D1 (**Figure 6C**).

**Figure 6:**
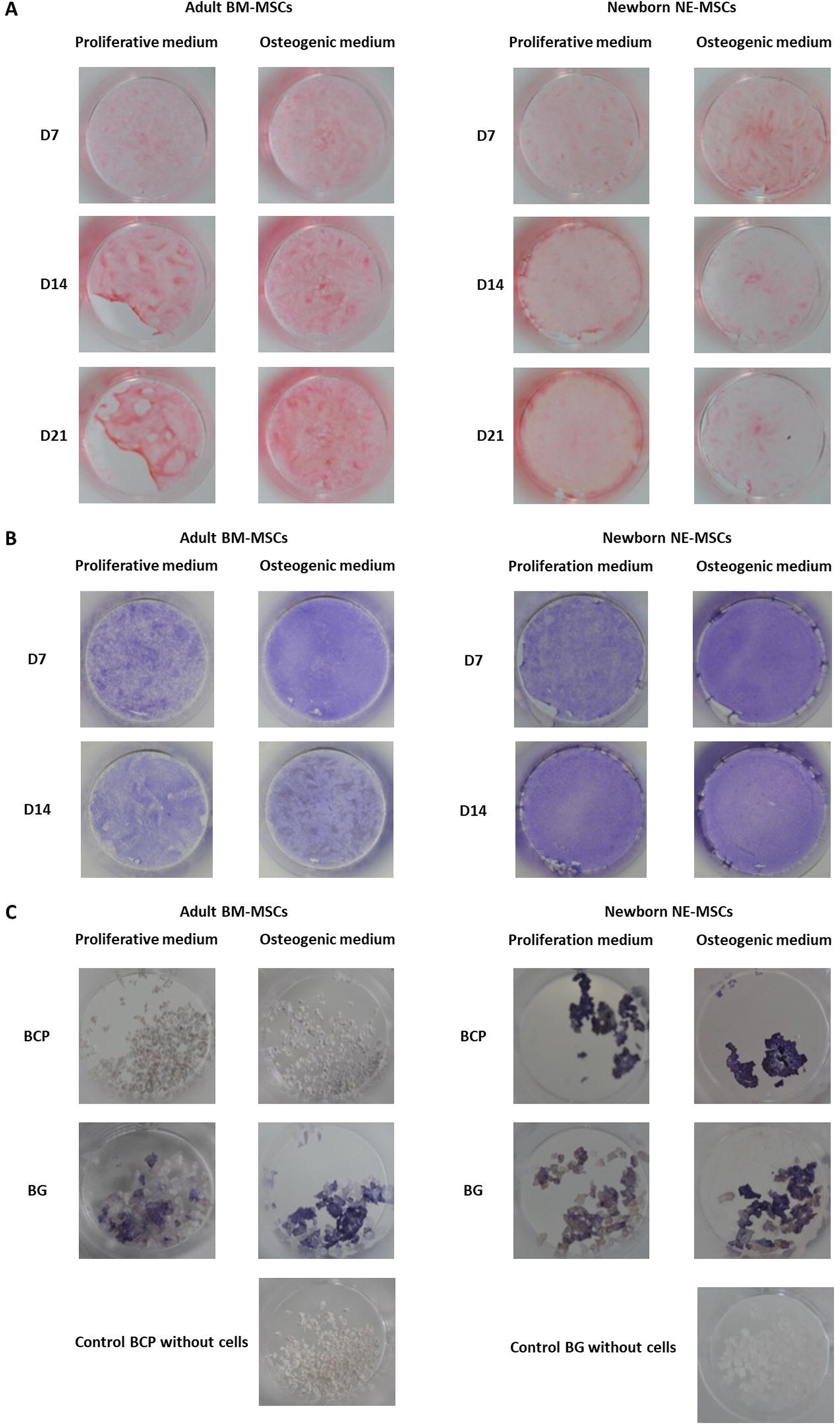
Quantification of alkaline phosphatase (ALP). For the purpose of comparison, the total amount of DNA, at D1 and D7, was quantified for both cell types BM-MSCs and NE-MSCs, in proliferative and osteogenic conditions (**A**). The total amount of *ALP* in each condition was assessed (**B**) and then compared to the amount of DNA (**C**). NE-MSCs express an important amount of *ALP* when grown on BG, in osteogenic conditions. For BM-MSCs, *ALP* expression is marginal except at D1 on BCP biomaterial.

### A differentiated expression of osteocyte-associated genes

Discrepancies between both MSCs further analyzed by measuring the expression of 8 bone-associated genes at D1 and D7 on both biomaterials (**Figure 7 and Supplementary Figure 4**). Normalization of gene expression performed using the amount expressed by BM-MSCs at D1 on BCP, as basal level. When cultivated on BG, NE-MSCs overexpress the genes coding for *BSP*, *OC* and *OP* (**Figures 7A-C**). Conversely, on the same biomaterial, BM-MSCs produce a higher number of transcripts coding for *RUNX2* (**Figure 7D**). No significant difference between the two groups noted for other genes (**Figures 8A, 8C, 8D**). *ALP* gene expression exhibited quite similar variations to ALP in culture supernatant, except for BCP at D1 (**Figure 8B**). No data are provided for BCP at D7 because a very small amount of RNA was extracted. This may be due to the failure of cell adhesion and proliferation on BCP, which can also explain the absence of correlation between ALP quantification and *ALP* gene expression at D1 on BCP. Indeed, reproducibility of experiment was affected by the absence of pre-incubation of BCP in culture medium prior to cell seeding.

**Figure 7:**
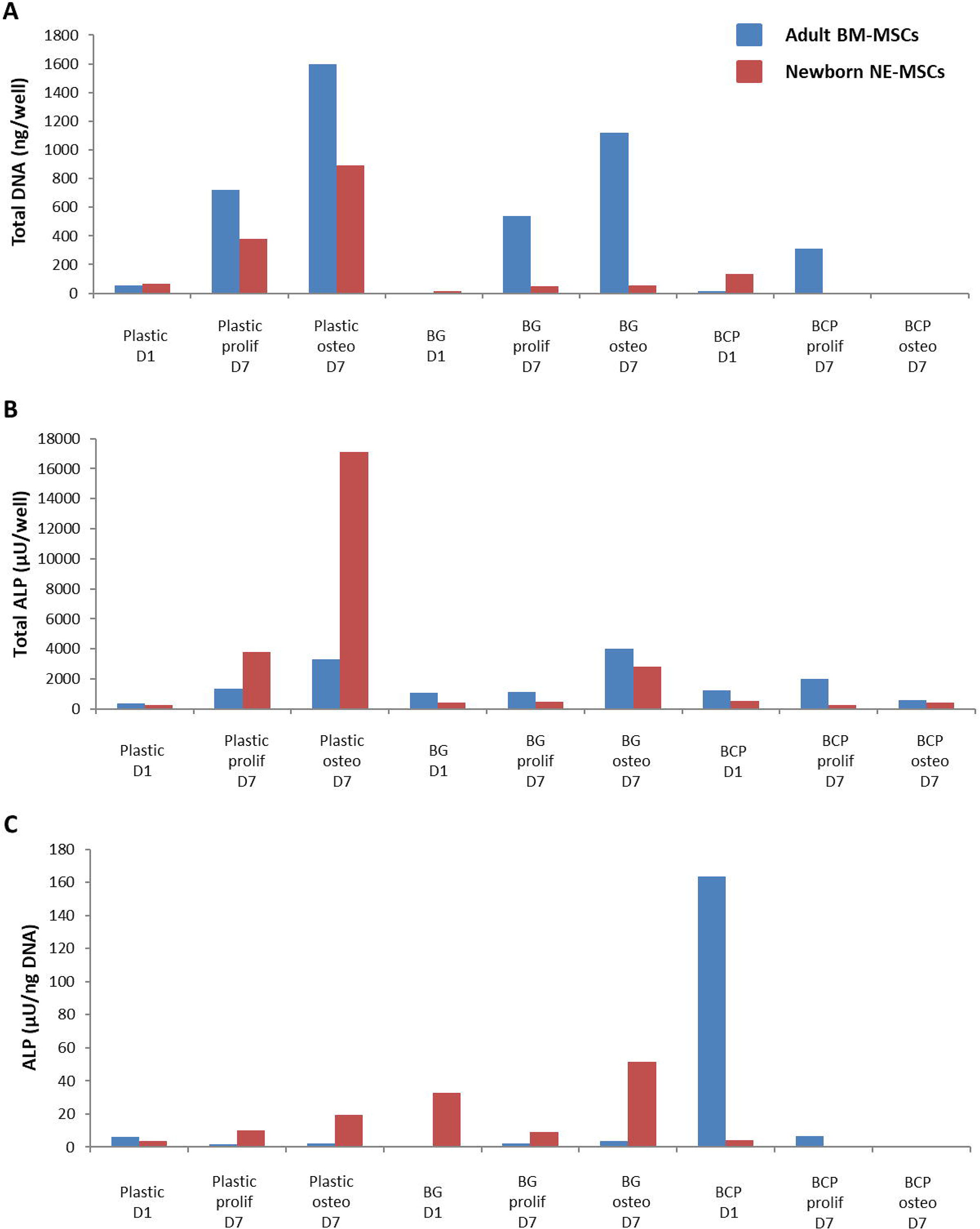
Comparative expression of ossification-related genes. Using RTqPCR, the expression of genes coding for *RUNX2*, *BSP*, *COL1A1*, *OC*, *BMP2*, *BMP4*, *ALP*, and *OP* was measured for each cell type on plastic, BG and BCP biomaterials. On BG biomaterial, NE-MSCs overexpress the genes coding for BSP, OC and OP (**A-C**) and BM-MSCs the gene coding for *RUNX2* (**D**).

**Figure 8:**
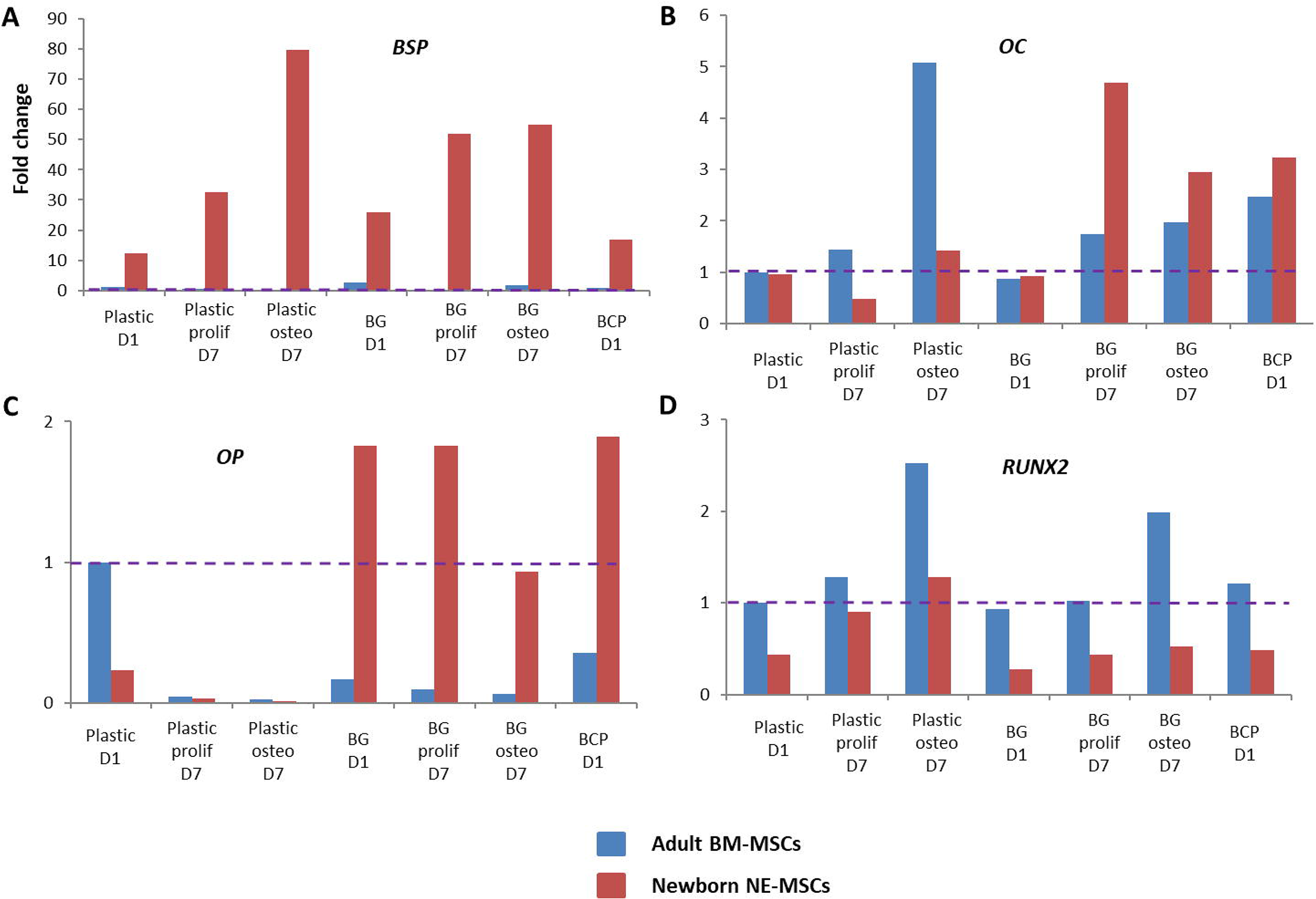
Expression of other ossification-related genes *COL1A1* (**A**), *ALP* (**B**)*, BMP2* (**C**), and *BMP4* (**D**) was measured by RTqPCR for each cell type on plastic, BG and BCP biomaterials. No data to D7 on BCP because the cells did not proliferate, and we obtained too little RNA.

### In vivo experiments

8 weeks after subcutaneous implantation in nude mice, none of the cell-free BG was found to harbor bone cells and vascularized fibrous tissue (**Figure 9A**). No bone formation was either observed on the same biomaterial loaded with BM-MSCs or NE-MSCs (**Figure 9C**). BM-MSCs gave rise to bone tissue when grafted on BCP (**Figure 9A-B**). Conversely, NE-MSCs failed to produce osteocytes (**Figures 9A-B**). No signs of systemic or local toxicity, no infections, and no behavioral changes were observed with both biomaterials with or without MSCs.

**Figure 9:**
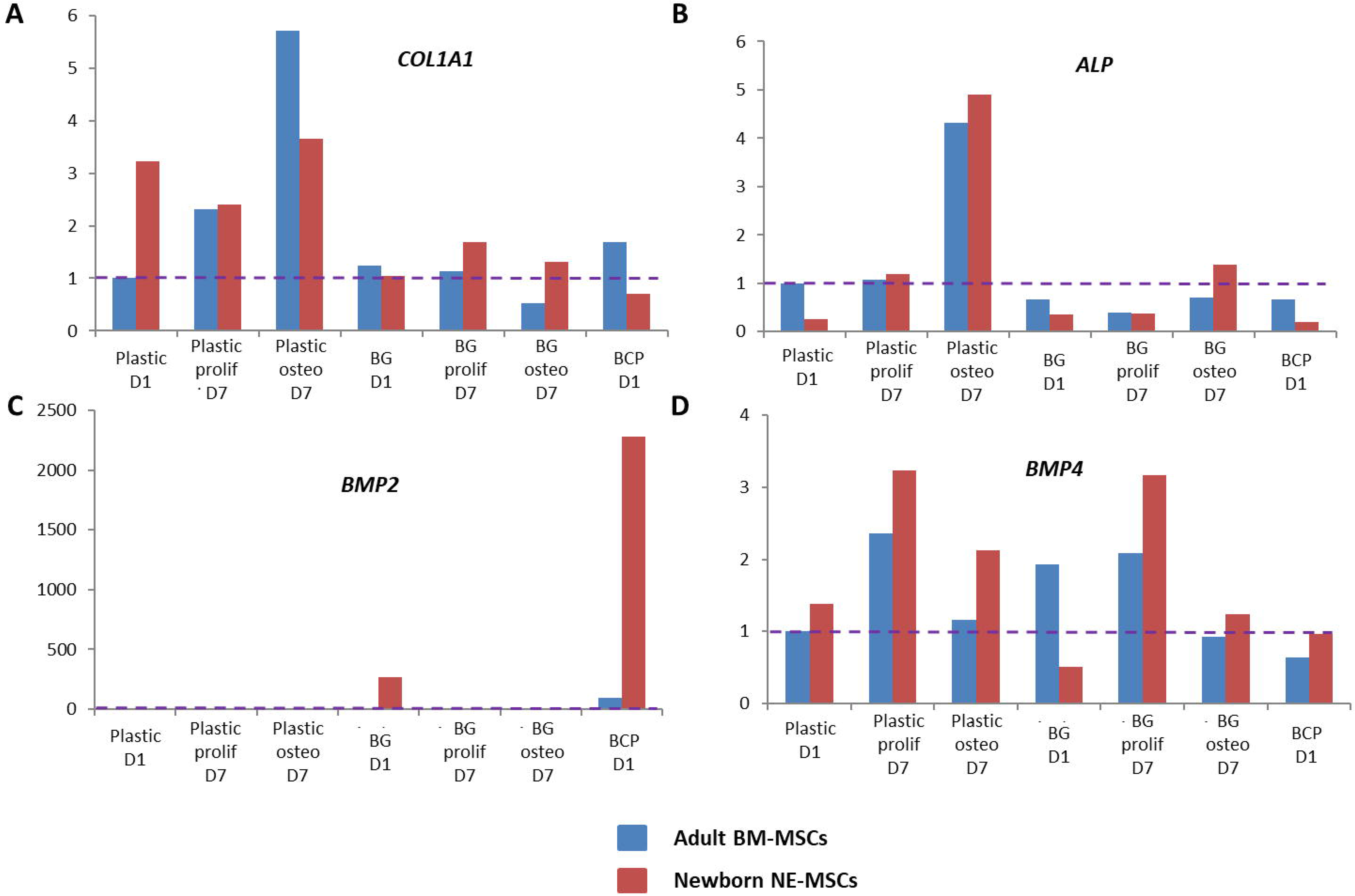
*In vivo* bone formation, 8 weeks after subcutaneous implantation in nude mice. (**A**) BM-MSCs give rise to osteocytes, stained with Masson trichrome (red arrows), but not NE-MSCs and cell-free BCP biomaterial. (**B**) The same finding is observed when the implants are analyzed with an electronic microscope (red arrows). (**C**) No bone formation is observed when cells were loaded in BG biomaterial. (**D**) Table summarizing the findings.

## Discussion

In this study, we compared the *in vivo* osteoinduction potential of hMSCs derived from either bone marrow or child nose associated with BCP or BG, two synthetic bone fillers currently used for their favorable bioactivity and osteoconductive properties^7,18,27^. Alone these biomaterials have insufficient osteoinductive properties to regenerate large bone defects. Combinations of hMSCs and BCP have been shown to induce *de novo* bone in ectopic sites and thus, have been widely studied *in vivo* as alternatives to autologous bone grafts^6,20^. On the opposite, combinations of MSCs and BG have been mainly studied *in vitro*,but poorly *in vivo*^24^. Although belonging to the same superfamily, BM-MSCs and NE-MSCs exhibit striking differences, *in vitro* and *in vivo*. For future clinical applications, the association of BM-MSCs with BCP biomaterial seems to be the most promising. Indeed, our *in vivo* experiments reveal that osteoinduction is only observed when BM-MSCs transplanted on BCP. Recently, Rodrigues and colleagues reported that allogenic adipose-derived MSCs associated with BG is i) biocompatible in the long term (3 months), ii) maintain their osteoinduction potential, and iii) is safe after subcutaneous implantation in immunocompetent *balb-c* mice. Indeed, a low spreading during cell adhesion was detected associated with an increase of HA depositions around the cells that looks like differentiated osteocytes. Any apparent local or systemic toxicity for organs, or strong immunogenic reactions were noted, except a vascularized dense capsule around the graft^**28**^.

Certain *in vivo* conditions could significatively impact the engraftment and the bone formation. In our study we chose to use nude mice to limit the immune response against the MSC allograft. The MSC allograft in Balb-c mice is possible, but the capsule formation surrounding the graft, probably due to the immune response in immunocompetent mice, could be a complication for implants^**28–30**^. The pH and the calcium concentration can also affect the cells. Here, we worked with 4 mM of calcium in culture medium. Maeno *et al.* showed that high concentration of calcium above 10 mM are cytotoxic to osteoblasts, but those below 8 mmol promote cell proliferation^**31**^. We also used medium supplemented with 5% GMP-grade human PL for MSC culture. It is now well established that animal derived products as FCS could significantly interact with phenotypical and functional characteristics of BM-MSCs^**25,26**^. Here, MSCs were grown with biomaterials 24 h before graft, and we implanted 2 × 10^6^ MSCs per site. In comparison, Rodrigues *et al.* used 2 × 10^4^ MSCs grown during 48 h on BG before implantation^**28**^.

## Materials and Methods

### Biomaterials

Two biomaterial scaffolds for hMSCs compared: macro/microporous BCP granules 0.5-1 mm in size (MBCP+^®^, Biomatlante, Vigneux de Betagne, France)^6,20^, and BG granules 0.5-1 mm in size (GlassBONETM, Noraker, Villeurbanne, France)^24^. BCP is a ceramic composed of HA/βTCP in a ratio of 20/80 by weight. BG 45S5 composed of 45 wt.% SiO_2_, 24.5 wt.% Na^2^O and 24.5 wt.% CaO.

### Donors

Bone marrow aspirates obtained from the iliac crest by standard puncture and aspiration of healthy human donors (21 and 26 years old), after receiving informed consent according to the Declaration of Helsinki. The project approved by the Ethical Committee of ULM University. BM-MSCs used in this study are of GMP grade, and expanded according to previously published protocols6. In brief, BM-MSCs isolated from heparinized bone marrow aspirates by seeding 50,000 white blood cells/cm^2^ on two-chamber CellStacks (Corning/VWR, Ulm, Germany) in α-MEM (Lonza, Basel, Switzerland) supplemented with 5% GMP-grade human platelet lysate (PL) (IKT, Ulm, Germany) in order to avoid animal products^25,26^. Cells cultured for 10 or 14 days with medium exchange twice per week. Cells detached and reseeded at a density of 4,000 BM-MSCs/cm^2^ on two-chamber CellStacks in α-MEM supplemented with 8% PL for a further 5 or 7 days.

Newborn NE-MSCs purified from the nasal mucosa, collected during a scheduled intervention for closing a cleft palate, under general anaesthesia. The procedure approved by an ethical committee (CPP, RCB: 2015-A00984-45). Two infants included in this pilot study. Biopsies sliced in small pieces, and incubated in 1.5 mL of collagenase (NB5, 1 U/mL, Nordmark Biochemicals) for 60 min at 37°C, before being mechanically dissociated. After digestion, 7 mL of serum containing culture medium added. After centrifugation, the cell pellet resuspended in DMEM/HAM supplemented with antibiotics (penicillin/gentamicin and fungizone), and Gibco^TM^ fetal calf serum (FCS) (Thermo Fisher Scientific, Waltham, Massachusetts, USA)on 25 cm^2^ plates. After 10 days, cells replated at a density of 4,000 cells/cm^2^ on 175 cm^2^ plates. At confluency, cells detached with Gibco^TM^ trypsin EDTA 0.05% (Thermo Fisher), and NE-MSCs stored in liquid nitrogen. Banking for scientific research performed anonymously, according to the rules of the Ministry of Higher Education and Research (number N°DC-2011-1331).

### Bioactive Glass pre-incubation

BG granules incubated in 1 ml of phosphate buffered saline (PBS) for 5 h, leading to a highly basic pH 10. In order to get a neutral pH, the BG granules (90 mg) incubated overnight in 1 mL of a calcium phosphate supersaturated solution (CPS). CPS contained 4 mM CaCl_2_.2H_2_O, and 2 mM of Na_2_HPO_4_.2H_2_O in 0.9% NaCl buffered at pH 7.4 with TRIS/HCl. After an overnight soaking in CPS, the pH of supernatant remained at 9. Based on these results, it was decided to seed the cells directly on the biomaterials without pre-incubation.

### Culture of MSCs with biomaterials

Culture of human BM-MSCs and NE-MSCs with biomaterials performed in Corning^®^ Costar^®^ ultra-low attachment 24 well plates (Merck KGaA, Darmstadt, Germany) with Gibco™ α-MEM (Thermo Fisher), supplemented with 8% human PL and 1% penicillin/streptomycin for 2-3 weeks (50 mg BCP or 90 mg BG per well).

### Adhesion of MSCs on biomaterials

Adhesion assessed using TM3000 scanning electron microscope (Hitachi, Krefeld, Germany), operating at an acceleration voltage of 5 kV, and imaged at a magnification of 50x-500x, after fixation and dehydration of the cells in ethanol.

### Viability and proliferation of MSCs

Cell viability on biomaterials evaluated at D2/7/14/21 by staining live cells with the fluorescent green stain calcein (1.25 μL/mL), and dead cells with fluorescent red ethidium homodimer-1 (1 μL/mL), according Invitrogen™ L-3224 kit (Thermo Fisher). Cell proliferation assessed for 21 days using alamarBlue^®^ assay (Thermo Fisher).

### hMSC characterization by flow cytometry

Cells stained with CD90-FITC, CD73-PE, CD105-PC7, CD45-APC-A750 antibodies (Beckman Coulter, Brea, California, USA), or corresponding isotype controls in matched concentration, 20 min at room temperature (RT) and protected from light. Cells washed in PBS without Ca^2+^ Mg^2+^ (Thermo Fisher), fixed, and permeabilized (IntraPrep Permeabilization Kit, Beckman Coulter). Intracellular staining then performed by indirect immunofluorescence method with an adapted dilution of Nestin antibody (Merck Millipore, Burlington, Massachusetts, USA), or its corresponding isotype control in matched concentration, and with an Alexa Fluor 647 goat anti-mouse IgG (H+L) secondary antibody (Thermo Fisher). After washing, cells analyzed with a NAVIOS flow cytometer (Beckman Coulter), and data files interprated using Kaluza software (Beckman Coulter).

### Tri-lineage differentiation capacity of expanded MSCs

**Osteogenic differentiation:** both BM-MSCs and NE-MSCs plated at the density of 5 × 10^3^ cells/cm^2^ in 24 well plates in basal media. After 1 day, MSC differentiation towards an osteogenic lineage induced with standard osteogenic supplements (10 mM *β*-glycerol-phosphate, 250 *μ*M ascorbic acid, and 100 nM dexamethasone). At D7/14/21, cells fixed with 4% paraformaldehyde (PFA), and mineralization detected by staining with a 40 mM alizarin red solution (pH 4.1-4.3).

**Adipogenic differentiation:** MSCs plated at the density of 2 × 10^4^ cells/cm^2^ in 24 well plates. Cells cultured until reaching 80% confluency in basal media and then induced towards adipogenic lineage with the StemPro™ Adipogenesis Differentiation kit (Thermo Fisher). After 14 or 21 days, cells fixed with 4% PFA, and adipocytes stained with Oil Red O solution in 2-propanol diluted to 60% using deionized water.

**Chondrogenic differentiation:** collected MSCs were loaded into 15 mL tubes (5 × 10^5^ cells/tube) in fresh basal media and centrifuged at 1,500 rpm during 5 min. After 24 h, the basal media removed, and StemPro™ Chondrogenesis Differentiation kit (Thermo Fisher) added to the cell pellet. At D14 or D21, cell pellets fixed in 4% PFA, and embedded in paraffin. Fixed pellets then cut with a microtome and stained with Alcian blue (Merck KGaA).

***Collagen production:*** visualized at D7/14/21 with Sirius red staining. Cells fixed with 4% PFA in PBS for 20 min at RT, then washed two times in PBS, and stained with 1 mL of 1 mg/mL of Direct red solution (Merck KGaA) in saturated aqueous solution of picric acid (Merck KGaA) for 1 h at RT.

**Extracellular alkaline phosphate (ALP)** qualitatively evaluated at D7/14/21 with Fast-Violet B Salt and Naphthol-AS-MX ALP staining (Merck KGaA). Cells fixed with a solution, containing two volumes of citric acid-sodium citrate (1.5 mol/L), and three volumes of acetone for 30 s at RT. Cells rinsed with deionized water and incubated with a staining solution for 30 min in the dark. 50 mL of staining solution contained 48 mL of distilled water, 12 mg of Fast-Violet B Salt (F1631, Merck KGaA), and 2 mL of Naphthol-AS-MX ALP solution (855, Merck KGaA).

### ALP quantification

At D1 and D7, MSCs were lysed using 0.1% Triton x-100, 5 mM Tris-HCL pH8 solution. After, three freeze/thaw cycles, the amount of double stranded DNA measured in the supernatants using a fluorescent Quant-iT Picogreen dsDNA Assay kit (Thermo Fisher), and the amount of ALP measured using SigmaFast™ p-Nitrophenyl phosphate (pNPP) tablets (Merck KGaA). A standard curve drawn with serial dilution of pNPP and a known quantity of ALP from bovine intestinal mucosa (Merck KGaA). Then, a known amount of pNPP added to each sample, prior to incubation during 30 min at 37°C. The amount of product (p-nitrophenol) determined by reading the absorbance at 405 nm on a microplate reader, and the amount of ALP quantified using the following equation:

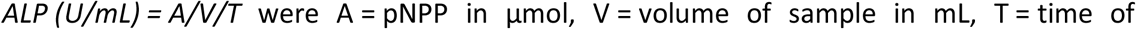

### Osteoblastic differentiation by real time quantitative PCR (RTqPCR)

Total RNA extracted from cultivated cells with DirectZol™ RNA MiniPrep (Zymo Research, Irvine, California, USA). The RNA samples (1 μg each) reverse transcribed with the maxima H minus First Strand cDNA Synthesis kit (Thermo Fisher) with oligo-dT primers in a final volume of 20 μL. The expression of each target gene normalized with glyceraldehyde 3-phosphate dehydrogenase (GAPDH) and beta-2-microglobulin (B2M). The 2-ΔΔCtmethod used to calculate relative expression levels. The relative gene expression normalized at D1 on plastic. Then the expression of the different bone coding genes assessed by RTqPCR.

### Primers

RUNX Family Transcription Factor 2 (*human RUNX2/F: gcctaggcgcatttcaga ; human RUNX2/R: gctcttcttactgagagtggaagg*), Bone sialoprotein (*human BSP/F: caatctgtgccactcactgc ; human BSP /R: cagtcttcattttggtgattgc*), Collagen Type I Alpha 1 Chain (*human COL1A1/F: ctggacctaaaggtgctgct ; human COL1A1/R: gctccagcctctccatcttt*), Osteocalcin (*human OC/F: tgagagccctcacactcctc ; human OC/R: ctggagaggagcagaactgg*), Bone morphogenetic protein 2 (*human BMP2/F: gttcggcctgaaacagagac ; human BMP2/R: ccaacctggtgtccaaaagt*), Bone morphogenetic protein 4 (*human BMP4/F: tcaagattggctgtcaagaatgatg ; human BMP4 /R: caggtatcaaactagcatggctcg*), Alkaline phosphatase (*human ALP/F: aacaccacccaggggaac ; human ALP/R: ggtcacaatgcccacagatt*), and Osteogenic protein (h*uman OP/F: gagggcttggttgtcagc ; human OP/R: caattctcatggtagtgagttttcc*).

### In vivo experiments

The *in vivo* experiment conducted according to the European regulation (Directive 2010/63/UE). The approval of the study obtained from the local ethical committee on animal experiment (number6575). Eighteen 7 week-old female nude mice used (NMRI nu/nu, Janvier Labs). 6 groups considered: BCP; BCP/BM-MSCs; BCP/NE-MSCs; BG; BG/BM-MSCs; BG/NE-MSCs on 50 mg of BCP or 90 mg of BG. No pre-treatment performed with BG granules. 2 × 10^6^ cells seeded on biomaterial granules and incubated overnight at 37°C with 5% CO2 prior to implantation. Two identical subcutaneous implantation per mouse performed. For statistics, 6 implants per group considered. Animals operated under general anaesthesia with Isoflurane (Abbvie, North Chicago, Illinois, USA). A centimetric skin incision performed on each side of mouse spine. Subcutaneous pockets filled with granules, embedded with or without MSCs. Wounds closed with non-resorbable suture 4.0 Filapeau (Péters Surgicals, Bobigny, France). Animals controlled every day post-surgery, to monitor the healing of the skin, and anomalies in behaviour. 8 weeks post-implantation, animals were euthanized, and implants fixed in 4% PFA.

### Histology and histomorphometry

BCP and BG samples dehydrated and embedded in poly-methyl-methacrylate resin (Merck KGaA). Sections performed with Leica SP1600 microtome (Wetzlar, Germany), and bone formation visualized by SEM. BCP samples decalcified in Decalc (Microm Microtech, Brignais, France), dehydrated in ascending series of alcohol, and embedded in paraffin. 4 μm-thin sections cut with a microtome in the middle of the implant. Sections stained by Masson trichrome in order to visualise newly formed bone tissue, granules, fibrosis, vascularization, and cells. Toxicity and biodegradation of biomaterials also determined.

### Statistics

Data processing and statistical analyses were performed with XLSTAT software (Microsoft, Redmond, Washington, USA). P-values less than 0.05 were considered significant. The results were expressed as mean +/− standard deviation (SD).

## List of abbreviations

MSCs: mesenchymal stromal stem cells
hMSCs: human mesenchymal stem cells
BM-MSCs: bone marrow derived MSCs
NE-MSCs: nasal ecto-mesenchymal stem cells
GMP: good manufacturing practice
BCP: biphasic calcium phosphate
HA: hydroxyapatite
βTCP: beta-tricalcium phosphate
BG: bioactive glass
PBS: phosphate buffered saline
CPS: calcium phosphate supersaturated solution
PFA: paraformaldehyde
RTqPCR: real time quantitative PCR
GAPDH: glyceraldehyde 3-phosphate dehydrogenase
B2M: beta-2-microglobulin
RUNX2: RUNX family transcription factor 2
BSP: bone salioprotein
COLA1: collagen A1
OC: osteocalcin
BMP2: bone morphogenic protein 2
BMP4: bone morphogenic protein 4
ALP: alkaline phosphatase
OP: osteopontin
RT: room temperature

## References

1. Logeart-Avramoglou D, Anagnostou F, Bizios R, Petite H. “Engineering bone: challenges and obstacles.” J Cell Mol Med. 2005;9(1):72–84.

2. Viateau V, Guillemin G, Calando Y, et al. “Induction of a barrier membrane to facilitate reconstruction of massive segmental diaphyseal bone defects: an ovine model.” Vet Surg. 2006;35(5):445–52.

3. Viateau V, Guillemin G, Bousson V et al. “Long-bone critical-size defects treated with tissue-engineered grafts: a study on sheep.” J Orthop Res. 2007;25(6):741–9.

4. Petite H, Viateau V, Bensaïd W et al. “Tissue-engineered bone regeneration. Nat Biotechnol.” 2000;18(9):959–63.

5. Pittenger MF, Mackay AM, Beck SC et al. “Multilineage potential of adult human mesenchymal stem cells.” Science. 1999;284(5411):143–7.

6. Brennan MÁ, Renaud A, Amiaud J et al. “Pre-clinical studies of bone regeneration with human bone marrow stromal cells and biphasic calcium phosphate.” Stem Cell Res Ther. 2014;5(5):114.

7. Stanovici J, Le Nail LR, Brennan MA et al. “Bone regeneration strategies with bone marrow stromal cells in orthopaedic surgery.” Curr Res Transl Med. 2016;64(2):83–90.

8. Brennan MA, Renaud A, Guilloton F et al. “Inferior In Vivo Osteogenesis and Superior Angiogenesis of Human Adipose-Derived Stem Cells Compared with Bone Marrow-Derived Stem Cells Cultured in Xeno-Free Conditions.” Stem Cells Transl Med. 2017;6(12):2160–2172. Erratum in: Stem Cells Transl Med. 2018;7(3):315.

9. Kaltschmidt B, Kaltschmidt C, Widera D. “Adult craniofacial stem cells: sources and relation to the neural crest.” Stem Cell Rev Rep. 2012;8(3):658–71.

10. Murrell W, Féron F, Wetzig A et al. “Multipotent stem cells from adult olfactory mucosa.” Dev Dyn. 2005;233(2):496–515.

11. Girard SD, Devéze A, Nivet E, Gepner B, Roman FS, Féron F. “Isolating nasal olfactory stem cells from rodents or humans. “ J Vis Exp. 2011;(54). pii: 2762.

12. Delorme B, Nivet E, Gaillard J et al. “The human nose harbors a niche of olfactory ectomesenchymal stem cells displaying neurogenic and osteogenic properties.” Stem Cells Dev. 2010;19(6):853–66.

13. Mankani MH, Kuznetsov SA, Fowler B, Kingman A, Robey PG. “In vivo bone formation by human bone marrow stromal cells: effect of carrier particle size and shape.” Biotechnol Bioeng. 2001;72(1):96–107.

14. Arinzeh TL, Tran T, Mcalary J, Daculsi G. “A comparative study of biphasic calcium phosphate ceramics for human mesenchymal stem-cell-induced bone formation.” Biomaterials. 2005;26(17):3631–8.

15. Mankani MH, Kuznetsov SA, Robey PG. “Formation of hematopoietic territories and bone by transplanted human bone marrow stromal cells requires a critical cell density.” Exp Hematol. 2007;35(6):995–1004.

16. Bruder SP, Kurth AA, Shea M, Hayes WC, Jaiswal N, Kadiyala S. “Bone regeneration by implantation of purified, culture-expanded human mesenchymal stem cells.” J Orthop Res. 1998;16(2):155–62.

17. Lebouvier A, Poignard A, Cavet M et al. “Development of a simple procedure for the treatment of femoral head osteonecrosis with intra-osseous injection of bone marrow mesenchymal stromal cells: study of their biodistribution in the early time points after injection.” Stem Cell Res Ther. 2015;6:68.

18. Gjerde C, Mustafa K, Hellem S et al. “Cell therapy induced regeneration of severely atrophied mandibular bone in a clinical trial.” Stem Cell Res Ther. 2018;9(1):213.

19. Gómez-Barrena E, Rosset P, Gebhard F et al. “Feasibility and safety of treating non-unions in tibia, femur and humerus with autologous, expanded, bone marrow-derived mesenchymal stromal cells associated with biphasic calcium phosphate biomaterials in a multicentric, non-comparative trial.” Biomaterials. 2019;196:100–108.

20. Gamblin AL, Brennan MA, Renaud A et al. “Bone tissue formation with human mesenchymal stem cells and biphasic calcium phosphate ceramics: the local implication of osteoclasts and macrophages.” Biomaterials. 2014;35(36):9660–7.

21. Rahaman MN, Day DE, Bal BS et al. “Bioactive glass in tissue engineering.” Acta Biomater. 2011;7(6):2355–73.

22. Jones JR. “Review of bioactive glass: from Hench to hybrids.” Acta Biomater. 2013;9(1):4457–86.

23. Kargozar S, Baino F, Hamzehlou S, Hill RG, Mozafari M. “Bioactive Glasses: Sprouting Angiogenesis in Tissue Engineering.” Trends Biotechnol. 2018;36(4):430–444.

24. Rizwan M, Hamdi M, Basirun WJ. “Bioglass^®^ 45S5-based composites for bone tissue engineering and functional applications.” J Biomed Mater Res A. 2017;105(11):3197–3223.

25. Ben Azouna N, Jenhani F, Regaya Z et al. “Phenotypical and functional characteristics of mesenchymal stem cells from bone marrow: comparison of culture using different media supplemented with human platelet lysate or fetal bovine serum.” Stem Cell Res Ther. 2012;3(1):6.

26. Leotot J, Coquelin L, Bodivit G et al. “Platelet lysate coating on scaffolds directly and indirectly enhances cell migration, improving bone and blood vessel formation.” Acta Biomater. 2013;9(5):6630–40.

27. Ebrahimi M, Botelho MG, Dorozhkin SV. “Biphasic calcium phosphates bioceramics (HA/TCP): Concept, physicochemical properties and the impact of standardization of study protocols in biomaterials research.” Mater Sci Eng C Mater Biol Appl. 2017;71:1293–1312.

28. Rodrigues C, Naasani LIS, Zanatelli C et al. “Bioglass 45S5: Structural characterization of short range order and analysis of biocompatibility with adipose-derived mesenchymal stromal cells in vitro and in vivo.” Mater Sci Eng C Mater Biol Appl. 2019;103:109781.

29. Velnar T, Bunc G, Klobucar R, Gradisnik L. “Biomaterials and host versus graft response: a short review.” Bosn J Basic Med Sci. 2016;16(2):82–90.

30. Chung L, Maestas DRJr, Housseau F, Elisseeff JH. “Key players in the immune response to biomaterial scaffolds for regenerative medicine.” Adv Drug Deliv Rev. 2017;114:184–192.

31. Maeno S, Niki Y, Matsumoto H et al. “The effect of calcium ion concentration on osteoblast viability, proliferation and differentiation in monolayer and 3D culture.” Biomaterials. 2005;26(23):4847–55.

